# SARS-CoV-2 Nsp1 regulates translation start site fidelity to promote infection

**DOI:** 10.1101/2023.07.05.547902

**Authors:** Ranen Aviner, Peter V Lidsky, Yinghong Xiao, Michel Tasseto, Lichao Zhang, Patrick L McAlpine, Joshua Elias, Judith Frydman, Raul Andino

## Abstract

A better mechanistic understanding of virus-host interactions can help reveal vulnerabilities and identify opportunities for therapeutic interventions. Of particular interest are essential interactions that enable production of viral proteins, as those could target an early step in the virus lifecycle. Here, we use subcellular proteomics, ribosome profiling analyses and reporter assays to detect changes in polysome composition and protein synthesis during SARS-CoV-2 (CoV2) infection. We identify specific translation factors and molecular chaperones whose inhibition impairs infectious particle production without major toxicity to the host. We find that CoV2 non-structural protein Nsp1 selectively enhances virus translation through functional interactions with initiation factor EIF1A. When EIF1A is depleted, more ribosomes initiate translation from an upstream CUG start codon, inhibiting translation of non-structural genes and reducing viral titers. Together, our work describes multiple dependencies of CoV2 on host biosynthetic networks and identifies druggable targets for potential antiviral development.

## Main

The process of viral replication relies heavily on the host’s translation machinery for the production of viral proteins and replication complexes, as well as the generation of infectious particle progeny. Such interactions are critical for successful infection, and most viruses have evolved to actively hijack and modulate their hosts’ protein synthesis machinery. This serves to both increase the efficiency of viral replication and prevent the production of antiviral factors^2^. Small molecule inhibitors of e.g. translation factors and molecular chaperones have shown antiviral effects across a wide range of viruses and animal models^3^, and some have entered clinical trials, suggesting that modulation of protein homeostasis (proteostasis) is a promising therapeutic approach. For CoV2, preclinical studies have reported antiviral efficacy for drugs targeting translation initiation factor 4A1 (EIF4A1) and elongation factor 1A (EEF1A)^4,5^. We previously demonstrated that proteomic analysis of ribosome-interacting proteins in cells infected with polio, Zika or dengue viruses can yield mechanistic insights into potential host targets for antiviral intervention^6^. Here, we use a similar approach to characterize CoV2 interactions with the cellular protein synthesis machinery. We separate translating and non-translating ribosomes from infected and control cells, and analyze their interacting proteins by liquid chromatography tandem mass-spectrometry (LC-MS/MS). We show that changes in ribosome interactors reflect biosynthetic requirements of CoV2. Specific translation factors and chaperones are recruited to support viral protein production, and their inhibition reduces infection. Interestingly, we find that CoV2 Nsp1 promotes efficient translation of CoV2 open reading frame 1 (Orf1), in a manner dependent on EIF1A, which prevents initiation from an alternative upstream start codon.

## Results

To identify host factors involved in CoV2 protein production, we infected Vero E6 cells with SARS-CoV-2 USA/WA1/2020 at a multiplicity of infection (MOI) of 5 plaque forming units (pfu) per cell, and performed subcellular fractionation coupled to mass-spectrometry analysis (Fig. 1a). Infected and control cells were lysed, fixed with formaldehyde, ultracentrifuged on 10-50% sucrose gradients, and fractionated with continuous monitoring of ribosomal RNA (rRNA) absorbance. As previously reported^7–10^, CoV2 infection reduced global translation, and a lower translation rate was maintained from 6 to 24 hours post-infection (hpi, Fig. 1b-c). A similar reduction was also measured in Calu3 cells (Extended Date Fig. 1a). Given that synthesis of viral proteins in Vero cells peaks between 10 and 24 hpi (Extended Date Fig. 1b)^11^, we chose 16 hpi for proteomic analysis. Lysates were subjected to sucrose gradient ultracentrifugation as above, and rRNA absorbance was used to determine how to pool fractions containing free 40S and 60S ribosomal subunits (free ribonucleoprotein complexes, “RNP”), 80S monosomes, and two polysome fractions corresponding to < and ≥ 6 ribosomes per polysome (“light” and “heavy” polysomes, respectively) (Fig. 1d). Based on measurements of protein concentration, the RNP fractions contained >90% of the protein mass present in the input lysates (Extended Data Fig. 1c). The protein content of each pooled fraction was then analyzed by label free LC-MS/MS (Supplementary Table 1). Overall, the composition of host proteins detected by MS was more variable between different fractions of the same gradient than between identical fractions of gradients from infected and uninfected cells (Fig. 1e). Given that gradient fractions separate cytoplasmic content based on size, this likely reflects common differences in protein composition between the monomeric proteome and larger protein assemblies. For example, higher correlations were measured between RNP fractions of infected and uninfected cells than RNP and any other fraction across the entire dataset (Fig. 1e). The abundance of ribosomal proteins in each pooled fraction, as quantified by MS (Fig. 1f-g and Extended Data Fig. 1d), was consistent with the observed rRNA absorbance profiles (Fig. 1d). Although our lysis conditions should disrupt the membrane-enclosed coronavirus replication-transcription complex (RTC)^12^, CoV2 RNA-dependent RNA-polymerase (RdRp, Nsp12) was still detected in the heavier fractions of the gradients (Extended Data Fig. 1e), possibly due to formaldehyde crosslinking. Nevertheless, MS estimates suggest that ribosomes outnumber RdRps in these fractions by a factor of about 500 to 1.

**Figure 1.**
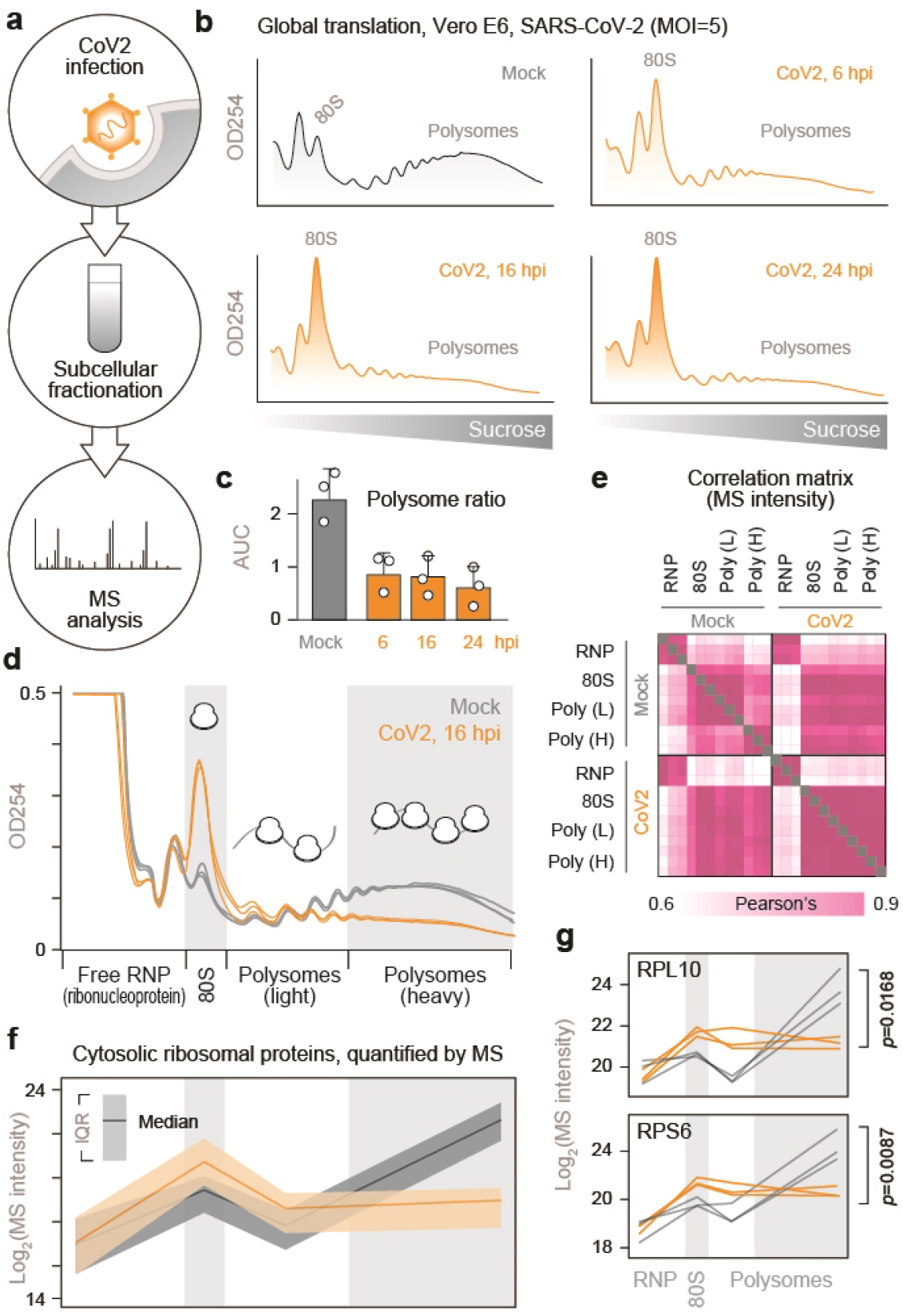
Subcellular proteomics of CoV2 infected cells. (**a**) To identify CoV2-mediated remodeling of host biosynthetic networks, we infected Vero E6 cells with SARS-CoV-2 USA/WA1/2020 at MOI=5, lysed them, fixed the clarified lysates with formaldehyde and fractionated on 10-50% sucrose gradients. Crosslinking was reversed and protein content was analyzed by liquid chromatography tandem mass-spectrometry (LC-MS/MS). (**b-c**) Global protein synthesis is persistently attenuated during infection. Cells infected as above were lysed, fixed with formaldehyde and fractionated on 10-50% sucrose gradients with continuous monitoring of rRNA absorbance (b). Ratio of polysomes to sub-polysomes, calculated as the area under the curve (AUC) of relevant fractions. Shown are means±SD of 3 independent replicates (c). (**d**) Cells were infected and fractionated as above, in triplicates, and protein content was extracted from fractions containing free small (40S) and large (60S) ribosomal subunits (free RNP); 80S monosomes; and two polysome fractions (“Light” and “Heavy”). Each line reflects a single replicate, and fractions pooled for MS are indicated at bottom. (e) Correlation matrix of all host proteins identified by MS in each of the pooled fractions from either CoV2-infected or control cells. (**f**) Median and interquartile range (IQR) of all cytosolic ribosomal proteins quantified by MS in each pooled fraction from all 3 replicates. (**g**) Line plots of individual ribosomal proteins quantified by MS in each pooled fraction. Each line represents a single replicate. P, two-tailed Student’s t-test p-value of differences in indicated protein abundance in heavy polysome fractions.

We next performed pairwise comparisons of non-ribosomal host proteins detected in each fraction of infected and control cells. CoV2 infection had little impact on the composition of free RNP and 80S fractions, but significantly altered the protein content of heavy polysome fractions (Fig. 2a). Many of the proteins recruited to heavy polysome fractions during CoV2 infection were previously shown to interact with either genomic or subgenomic CoV2 RNA^13^ (Fig. 2a, pink). Most of these enriched proteins are involved in RNA metabolism, protein synthesis and maturation, including RNA splicing and transport, translation, protein folding, proteasomal degradation and antigen presentation (Fig. 2b and Supplementary Table 2). As previously reported, these co-translational interactions are mediated by both viral RNA and nascent polypeptide chains^6^. Individual examples include splicing factors RTCB, NONO and PPP1R8—three of the top 15 hits in a CRISPR screen for essential CoV2 host factors^13^, as well as SRSF5 and HNRNPD, previously shown to affect RNA splicing, translation and stability in other viruses^14,15^ (Fig. 2c and Extended Data Fig. 2). Additional host factors recruited to CoV2 polysomes include subunits of the 26S proteasome (Fig. 2c), potentially reflecting increased co-translational degradation of nascent polypeptide chains that are sensed as aberrant by host protein quality controls^16^. Other components of the ubiquitin-proteasome system were also recruited, such as HSPA5/BiP (the ER-resident Hsp70) and DNAJC3/Erdj6, which mediate interactions with proteasomes to facilitate co-translational degradation of polypeptides on the ER (reviewed in ^17^); and TRIM25, an E3 ligase of the antiviral ISG15 conjugation system, which targets nascent viral proteins^18^ (Extended Data Fig. 2).

**Figure 2.**
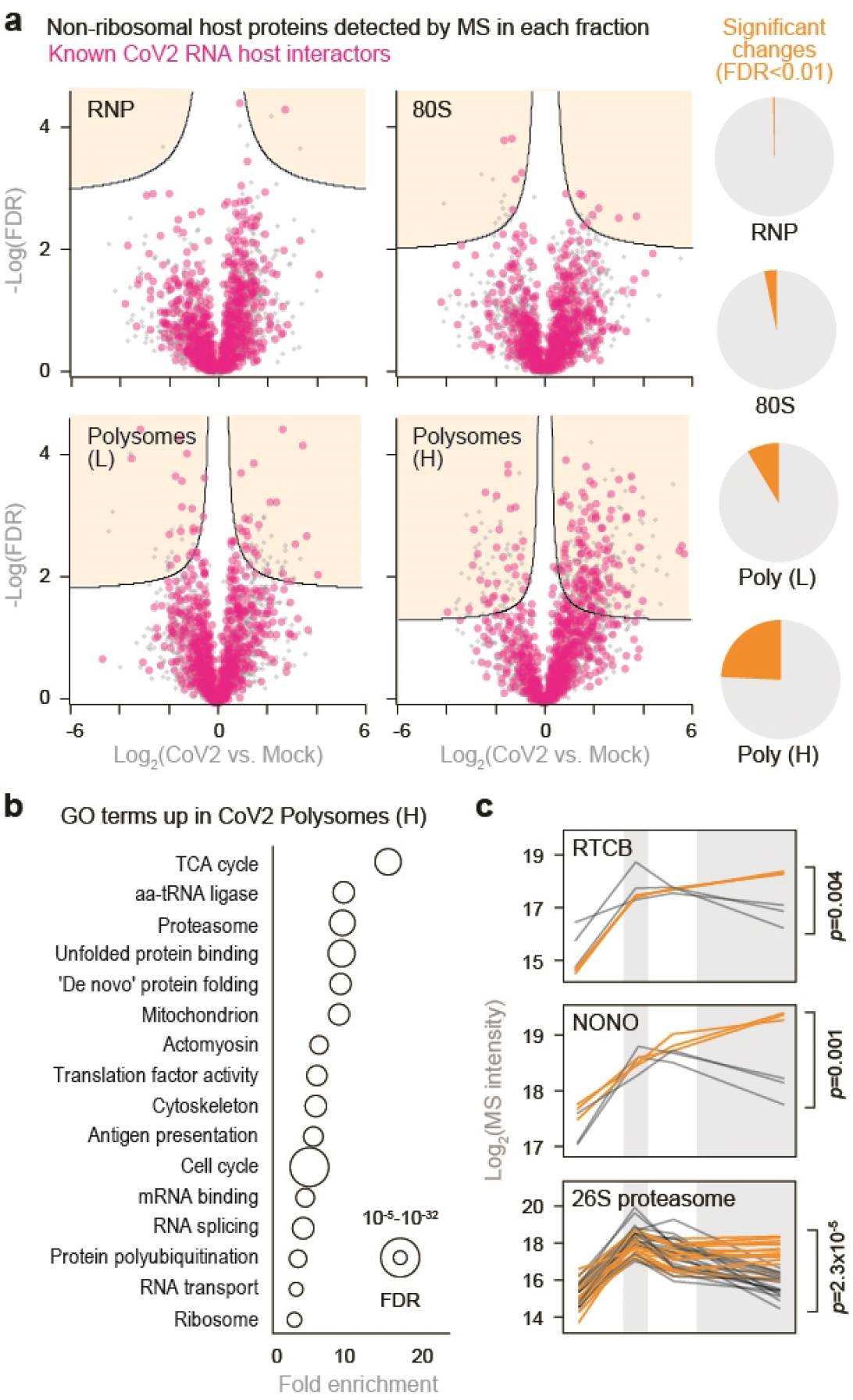
CoV2 infection remodels host biosynthetic complexes. (**a**) Pairwise comparisons of differences in individual protein abundance upon CoV2 infection of Vero E6 cells, per pooled fraction, as quantified by MS. Right, proportion of proteins showing statistically significant differences (FDR<0.05, S0=0.1) between infected and control cells. (**b**) Gene Ontology terms enriched in heavy polysome fractions from infected versus control cells. (**c**) Line plots of individual proteins quantified by MS in each fraction. Each line represents a single replicate. P, two-tailed Student’s t-test p-value.

To determine whether knowledge of polysome remodeling during CoV2 infection may provide information on virus vulnerabilities that can be targeted for antiviral interventions, we initially focused on molecular chaperones and other components of proteostasis, which are critical to the successful production of functional virus proteins^3^. We infected Vero cells with CoV2 at MOI=0.5 in the presence of either DMSO or drugs that target some of the host factors recruited to heavy polysome fractions during infection (Fig. 3a). These include Juglone, an inhibitor of prolyl isomerase of the parvulin family; 16F16, a protein disulfide isomerase inhibitor; JG40 and JG345, which inhibit Hsp70 chaperones; and Nimbolide, which inhibits RNF114 E3 ligase. Remdesivir, a known inhibitor of the viral RdRp, was used as a positive antiviral control. At concentrations not toxic to host cells, all drugs inhibited CoV2 infection (Fig. 3b and Extended Data Fig. 3a-b), suggesting that the virus is hyper-dependent on these targeted functions. Furthermore, combined inhibition of disulfide isomerases and Hsp70, but not disulfide isomerases and remdesivir or any other combination, had synergistic antiviral effects (Fig. 3c). Together, these observations confirm that the information generated in the context of these polysome analyses can be used to develop effective antiviral interventions targeting key steps in the viral life cycle.

**Figure 3.**
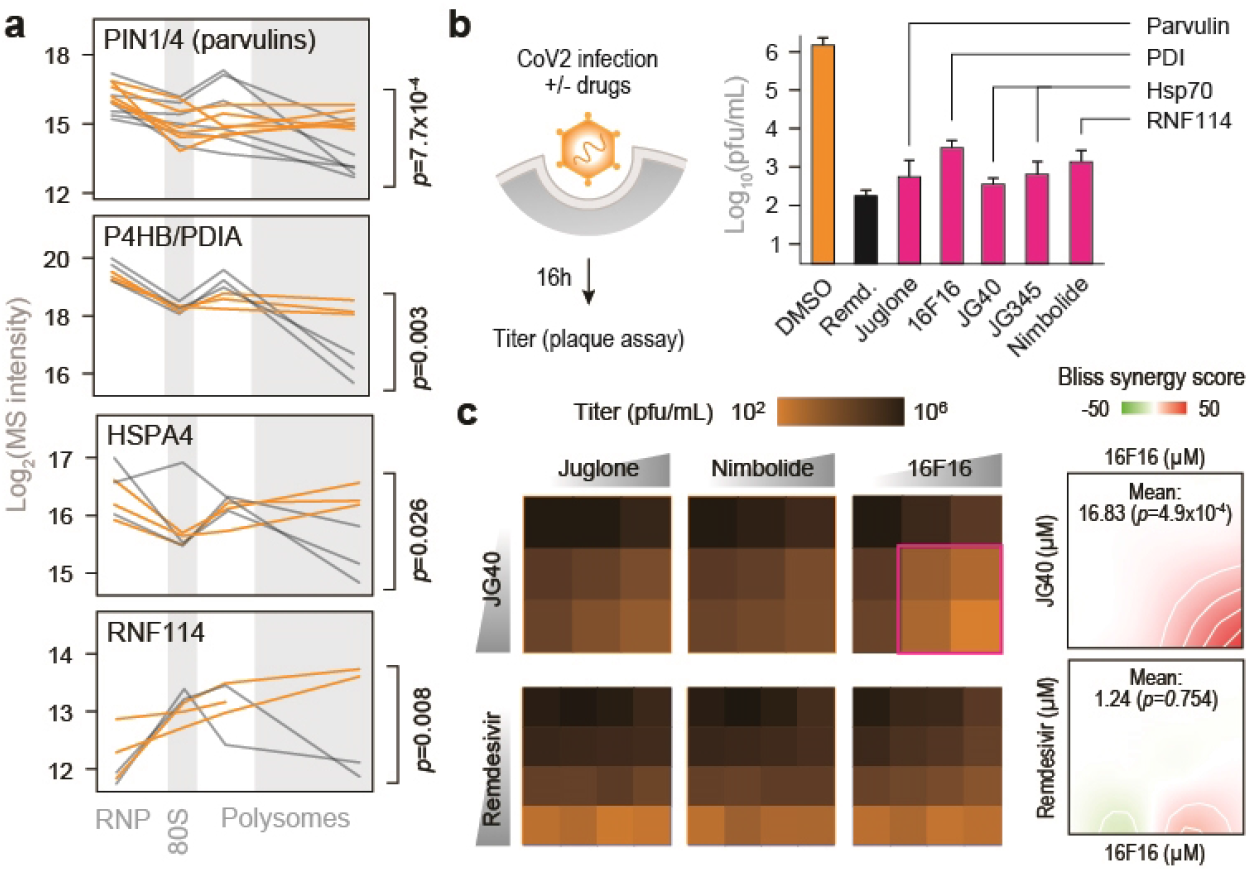
Remodeling of biosynthetic complexes reveals druggable host targets for antiviral therapies. (**a**) Line plots of individual proteostasis factors quantified by MS in each fraction. Each line represents a single replicate. P, two-tailed Student’s t-test p-value. (**b**) Cells were infected with CoV2 at MOI=0.5. Drugs were added at the start of infection, and titers were determined by plaque assays at 16 hours post-infection. Shown are means±SD of 3 independent replicates. Remdesivir, 5 µM; Juglone, 4 µM; 16F16, 2 µM; JG40, 5 µM; JG345, 5 µM; Nimbolide, 1 µM. (**c**) Cells were infected as above and treated with the indicated drug combinations. JG40, 0, 2.5, 5 µM; Remdesivir, 0, 2.5, 5, 10 µM; Juglone, 0, 0.25, 0.5, 1 µM; Nimbolide, 0, 0.25, 0.5, 1 µM; 16F16 0, 1, 2, 4 µM. Shown are means of 3 independent replicates. (**d**) Bliss synergy score for combined treatment with 16F16 and either JG40 or remdesivir. Higher values indicate synergistic effects.

### Translation initiation on CoV2 gRNA is inefficient

Next, we examined whether CoV2 infection affects the core components of translating ribosomes. Consistent with a global translation shutoff, heavy polysome fractions from infected cells contained fewer elongating ribosomes, reflected by lower abundance of ribosomal proteins and the two major elongation factors, EEF1A and EEF2 (Fig. 4a). In contrast, multiple translation initiation factors were enriched in heavy polysome fractions (Fig. 4a), but not other fractions (Extended Data Fig. 4a), upon CoV2 infection. These included EIF1, 1A, 3A, 4A1, 4B and DDX3 (Fig. 4a-b and Extended Data Fig. 4b). By reanalyzing previously published datasets^13^, we found that translation initiation factors are also some of the most abundant CoV2 RNA interactors in Vero cells (Fig. 4c). Furthermore, viral RNAs are associated with more 40S than 60S ribosomal subunits (Fig. 4c, inset), likely representing pre-initiation complexes prior to large subunit joining. Similar observations were also made based on a separate study of CoV2 RNA interactors in Huh7 cells^19^ (Extended Data Fig. 4c). Interestingly, our sucrose-fractionated rRNA absorbance profiles of CoV2-infected cells show lower abundance of free 40S subunits in both Vero (Fig. 4d) and Calu3 (Extended Data Fig. 4d). In addition, more 40S subunits are found by MS in the heavy polysome fractions of infected Vero cells (Fig. 4e). These observations suggest that the assembly of 80S initiating ribosomes is inefficient, and perhaps an important limiting step during CoV2 infection.

**Figure 4.**
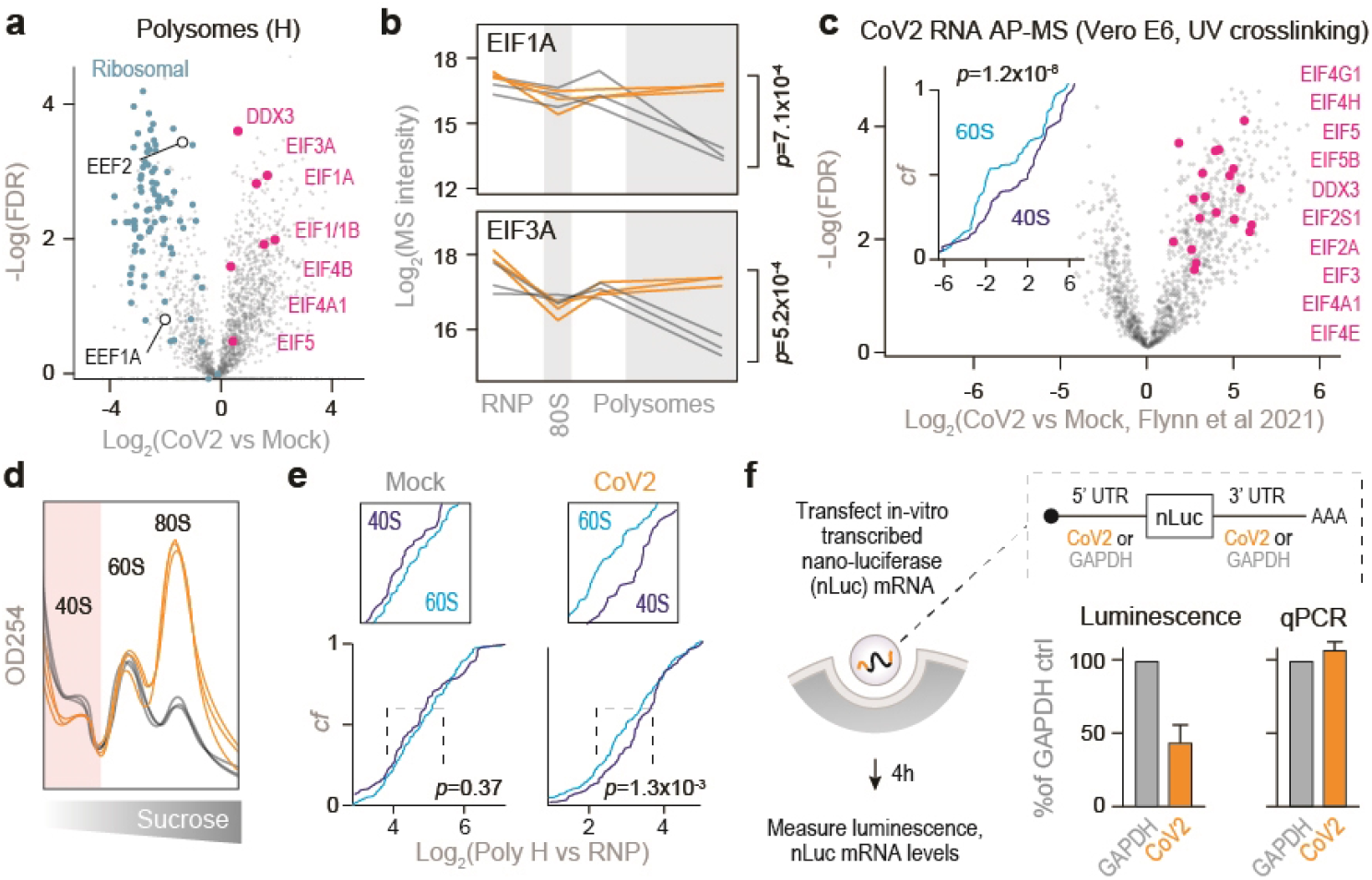
Inefficient translation initiation on CoV2 gRNA. (**a**) Pairwise comparisons of individual protein abundance in the heavy polysome fractions. Ribosomal proteins in blue, translation initiation factors in pink, and translation elongation factors in white circles. (**b**) Line plots of individual translation initiation factors quantified by MS in each fraction. Each line represents a single replicate. P, two-tailed Student’s t-test p-value. (**c**) CoV2 RNA interactome is enriched for translation initiation factors during infection. Shown are pairwise comparisons of individual host protein abundance, quantified by MS, that specifically interact with either genomic or subgenomic CoV2 RNA. Inset, cumulative distribution plots of 40S and 60S ribosomal protein interaction with CoV2 RNA. *P*, Mann-Whitney p-value. (**d**) rRNA absorbance profiles from Fig. 1D, showing lower abundance of free 40S subunits during CoV2 infection. (**e**) Heavy polysome fractions contain more 40S ribosomal proteins in infected cells. Shown are cumulative distribution plots of 40S and 60S ribosomal proteins in heavy polysome fractions from infected and uninfected cells, across three replicates. *P*, Mann-Whitney p-value. (**f**) Translation initiation from CoV2 5’ untranslated region (UTR) is less efficient than GAPDH 5’UTR. mRNA encoding for nano-luciferase (nLuc) flanked by 5’ and 3’UTRs of either CoV2 or GAPDH was transcribed in vitro, capped/polyadenylated, and transfected into Vero E6 cells. At 4 hours post-transfection, luminescence was measured in parallel with qPCR using oligonucleotides specific to nLuc. Shown are means±SD of 3 independent replicates.

To test the rate of translation initiation on viral RNA, we in-vitro transcribed and capped a polyadenylated nano-luciferase (nLuc) mRNA flanked by 5’ and 3’ untranslated regions (UTR) of either GAPDH or CoV2 gRNA^10^. As the two mRNA variants encode for an identical protein sequence, differences in luminescence should reflect the translation initiation efficiency of each set of UTRs. We chose RNA transfections to overcome potential confounding factors associated with plasmid DNA transfection e.g. transcription, splicing, nuclear modifications and export^20^. At 4h post-transfection in Vero cells, we measured nLuc luminescence and mRNA levels. Despite similar levels of intracellular mRNA for both reporters, CoV2 UTRs generated about 2-fold less nLuc luminescence than GAPDH UTRs (Fig. 4f). Thus, it appears that translation initiation on CoV2 RNA is inefficient, due to regulatory sequence or structural elements present in its UTRs. This is consistent with previously measurements using ribosome profiling^7^.

### Nsp1 promotes translation initiation on CoV2 gRNA

Coronavirus Nsp1 is known to inhibit global translation by binding to 40S subunits and interfering with initiation^8,9^. CoV2 gRNA escapes this repression, likely by displacing Nsp1 from the mRNA entry channel of the ribosome^21^. To determine whether Nsp1 is displaced from polysomes, we compared its sedimentation pattern to those of other viral proteins translated from the same Orf1 polyprotein. As expected, Nsp1 was detected mostly in the free RNP and 80S fractions of infected cells, while almost all other viral proteins were enriched in the heavier fractions of the gradients (Fig. 5a and Extended Data Fig. 5a). Nevertheless, Nsp1 was still detected at significant levels in polysome fractions. To determine how individual viral proteins affect translation initiation on CoV2 gRNA, we repeated the mRNA reporter assays in the presence of individually expressed viral proteins, and measured luminescence and association of nLuc mRNA with translating ribosomes (Fig. 5b). Most viral proteins had negligible effects on translation initiation from either GAPDH or CoV2 UTRs. In contrast, Nsp1 reduced translation driven by GAPDH UTRs and increased translation driven by CoV2 UTRs (Fig. 5c). To confirm that these effects are directly related to translation initiation, we expressed either Nsp1 or GFP, transfected the same translation reporters, and analyzed polysome association of nLuc mRNA using RT-qPCR analysis of sucrose gradient fractions. Compared to GFP control, Nsp1 reduced global translation rates, shifting GAPDH-nLuc mRNA away from heavy polysomes (Fig. 5d and Extended Data Fig. 5b). In contrast, Nsp1 recruited CoV2-nLuc mRNA to polysomes despite a similar reduction in global translation (Fig. 5d and Extended Data Fig. 5b).

**Figure 5.**
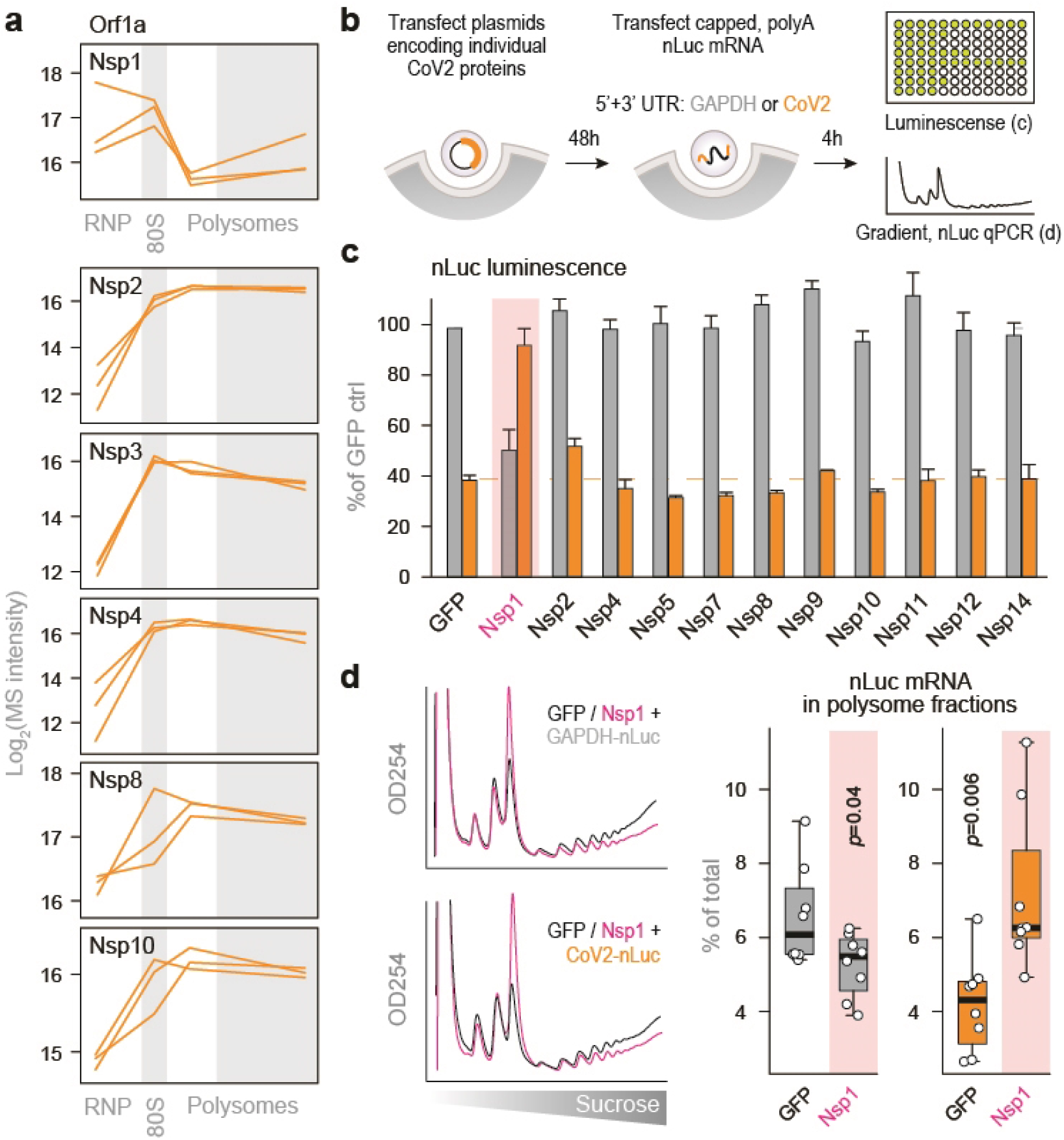
Nsp1 promotes translation initiation on CoV2 gRNA. (**a**) Line plots of individual viral proteins quantified by MS in each fraction. Each line represents a single replicate. (**b**) Vero E6 cells were transfected with plasmids encoding for individual CoV2 proteins. At 48 hours post-transfection, cells were transfected with nLuc mRNA flanked by either CoV2 or GAPDH UTRs. At 4 hours post-second transfection, cells were subjected to either luminescence measurements or sucrose gradients coupled to qPCR of nLuc mRNA. (**c**) nLuc luminescence. Shown are means±SD of 3 independent replicates. (**d**) Cells transfected as above, with either GFP or Nsp1 followed by GAPDH-nLuc or CoV2-nLuc mRNA, were lysed and fractionated on 10-50% sucrose gradients with continuous monitoring of rRNA. The content of nLuc mRNA in each fraction was determined by qPCR. Left, rRNA absorbance profiles. Right, percent of GAPDH-nLuc or CoV2-nLuc mRNA found in polysome fractions of sucrose gradients. Shown are qPCR measurements of 4 polysome fractions from 2 independent gradients. P, two-tailed Student’s t-test p-value.

### EIF1A guides start site selection and mediates the effects of Nsp1 on CoV2 gRNA translation

To learn more about the role of Nsp1 in translation initiation of CoV2 gRNA, we investigated potential links between Nsp1 and translation factors. Re-analysis of two datasets using proximity labeling coupled to proteomic analysis of Nsp1 interactors^22,23^ revealed that wild-type Nsp1, but not Nsp1 deficient for ribosome binding, selectively interacts with a subset of translation initiation factors (Fig. 6a), some of which were also enriched in heavy polysome fractions from CoV2-infected cells (Fig. 4b). This suggested that Nsp1 may preferentially interact with or stabilize 48S pre-initiation complexes with specific composition or function, as supported by cryo-EM studies of Nsp1-bound ribosomes^9^.

**Figure 6.**
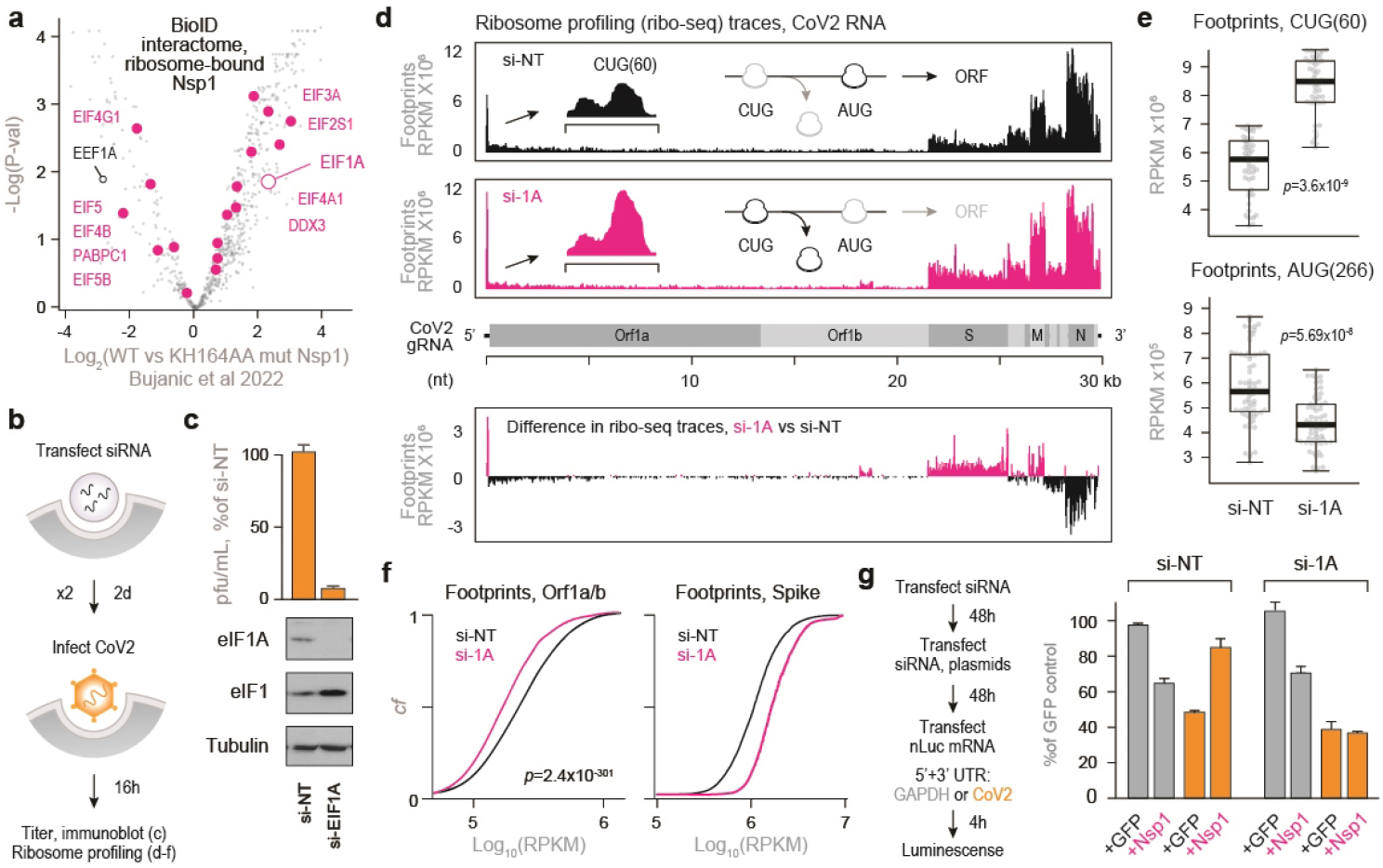
Nsp1 promotes accurate start codon usage through eIF1A. (**a**) Nsp1 interacts with translation initiation complexes enriched for specific initiation factors. Pairwise comparisons of individual protein abundance that, as quantified by MS, specifically interact with Nsp1 using a bioID proximity labeling approach. (**b**) Vero E6 cells were transfected with either non-targeting or eIF1A and eIF1-targeting siRNA. At 48h the same transfection was repeated. At 48h after the second transfection, cells were infected with CoV2 at MOI of either 0.1 (c) or 5 (d-f). Titer was determined by plaque assays. (**c**) Shown are means±SD of 3 independent replicates. Bottom, immunoblot of whole cell lysates transfected with the indicates siRNAs. (**d**) Infected cells were subjected to ribosome profiling analysis. Top, ribosome footprints on CoV2 vRNA. Insets, ribosome footprints on nucleotides 20-80 of the genomic vRNA. Representative of 2 independent replicates. Bottom, difference in ribosome occupancy at each codon between cells pre-transfected with either si-eIF1A or si-NT. (**e**) Ribosome occupancy at nucleotides 48-78 (top, upstream CUG codon) and 254-284 (bottom, main AUG). Summary of two independent replicates. P, Wilcoxon ranked-sum p-value. (**f**) Cumulative fraction plots of ribosome footprints on Orf1a/b (left) and Spike (right) vRNA. Summary of two independent replicates. P, Wilcoxon ranked-sum p-value. (**g**) Vero E6 cells were transfected with either si-eIF1A or si-NT. At 48h, cells were retransfected with same siRNA and plasmids encoding for either GFP or Nsp1. 48h later, cells were transfected again with nLuc mRNA flanked with either CoV2 or GAPDH UTRs. Luminescence was measured at 4h. Shown are means±SD of 3 independent replicates.

Ribosome profiling analyses have shown that, as ribosomes initiate translation on CoV2 gRNA, they accumulate on a CUG codon upstream of the main translation start site^24,25^. This conserved CUG at position 60 of the gRNA is out of frame compared to the main AUG at position 266, and predicted to suppress translation of Orf1^25^. We searched through our datasets for initiation factors that are both associated with Nsp1 and recruited to heavy polysome fractions during infection. This highlighted EIF1A (Fig. 6a, 4b), which is known to restrict translation initiation from non-AUG start codons^26^. To test the importance of EIF1A in CoV2 infection, we transfected Vero cells with either non-targeting or anti-EIF1A siRNA, infected with CoV2, and measured virus production by plaque assays and viral translation by ribosome profiling (Fig. 6B). Knockdown of EIF1A increased the levels of EIF1 through a known negative feedback loop^26^ and reduced CoV2 titers by about 10 fold (Fig. 6c). Ribosome profiling analysis confirmed that knockdown of EIF1A increases ribosome accumulation on the upstream CUG and decreases ribosome accumulation on the main AUG of Orf1 (Fig. 6d-e). This was associated with lower ribosome occupancy on the coding region of Orf1, but higher occupancy on the coding region of Spike (Fig. 6f). This may be explained by differences in UTR sequences that are introduced by subgenomic transcription downstream of CUG(60)^27^. In Spike subgenomic RNA, the upstream CUG(60) is in frame with the main AUG, and a CUG-to-CCG mutation reduces its translation^25^.

Finally, we examined whether EIF1A is required for Nsp1-mediated enhancement of CoV2 gRNA translation. We sequentially transfected Vero cells with siRNAs, GFP or Nsp1 plasmids, and GAPDH-or CoV2-nLuc mRNA. In control cells, Nsp1 reduced translation from GAPDH UTR and stimulated translation from CoV2 UTR. However, in the absence of EIF1A, Nsp1 still reduced translation from GAPDH UTR but failed to stimulate translation from CoV2 UTR (Fig. 6g). Taken together, we conclude the EIF1A is required for accurate translation start site selection on CoV2 gRNA, and that Nsp1 depends on EIF1A to promote efficient translation of CoV2 gRNA.

## Discussion

Despite intense research efforts, much is still unknown about the complex interactions between CoV2 and its host cell. Protein-protein and protein-RNA interactions have been characterized using various methods^4,13,19,28^, but their functional significance remains poorly understood, and more work is needed to determine whether they reflect druggable viral dependencies. Several lines of evidence suggest that co-translational CoV2 interactors may be a particularly promising class of targets for host-directed antivirals, due to the unique challenges associated with its protein synthesis. The viral RNAs are highly structured^28^ and harbor multiple overlapping open reading frames^24^, two of which encode for long multifunctional polyproteins. Orf1ab, for example, is synthesized as a single protein of 7,096 amino acids—more than 10 times the average length of a human protein^29^—and then processed into 16 individual subunits, each with its own unique structure, function, and interaction networks^4,30^. The recruitment of diverse protein translation, folding and degradation factors to CoV2 polysomes, as found in this study, as well as identification of selective antiviral effects of several translation inhibitors^4,5^, further extends our knowledge of such unique biosynthetic dependencies.

Nevertheless, data on CoV2 translation efficiency remains ambiguous. Some reporter assays suggest that translation of all CoV2 RNA species is highly efficient, or at a minimum similar to that of cellular mRNAs encoding for housekeeping proteins^10,21,31^. In contrast, ribosome profiling and other sequencing- and immunofluorescence-based studies show that viral translation efficiency is lower than that of cellular mRNAs^7,32^, although some measurements may be biased by the presence of positive-strand viral RNA in replication, transcription and packaging complexes. Furthermore, a detailed mutagenesis analysis of CoV2 5’UTR identified multiple sequence and structure elements that reduce translation initiation from the main AUG^33^. Such discrepancies may also be explained by differences in experimental design, which ranges from in-vitro translation to DNA or RNA transfection to studies of translation in infected cells.

Furthermore, multiple reports have argued that Nsp1 inhibits translation initiation of cellular mRNAs but does not affect that of CoV2 gRNA^7–10,21^. However, at low concentrations, Nsp1 was shown to promote gRNA translation^22^. Our results support the latter model: in our hands, Nsp1 suppressed translation from GAPDH UTRs and promoted that of CoV2 gRNA UTRs. We find that this stimulatory function of Nsp1 depends on a specific initiation factor, EIF1A. The C-terminal part of Nsp1 binds the 40S ribosome in close proximity to EIF1A^9^, which plays a role in start site selection^26^. Furthermore, EIF1A is present in Nsp1 pulldowns^9,22,23^, confirming the two proteins co-occupy pre-initiation complexes. It is tempting to speculate that Nsp1 stabilizes EIF1A binding or otherwise affects its interactions with the ribosome decoding center, although additional work is needed to elucidate the specifics of the mechanism. Nevertheless, in the absence of EIF1A, Nsp1 fails to promote gRNA translation and ribosomes accumulate on an inhibitory upstream CUG(60), one of several translation start sites in CoV2 5UTR^24^.

Upstream open reading frames (uORFs) are common translation repressors^34^ that shape the proteome during cellular stress^35^, and AUG-or CUG-initiated uORFs are found in most known coronaviruses^36^. A conserved uORF in mouse hepatitis virus (MHV) suppresses translation of Orf1 and plays a nonessential role in viral replication in cell culture, but mutations disrupting its function are quickly reversed upon passaging^36^. Similarly, a conserved uORF in enteroviruses is dispensable for replication in cell culture but promotes infection in gut epithelial cells^37^. These observations suggest that uORFs play important yet undefined roles in permissibility of specific cell types to viral infection.

Together, our findings reveal a functional interplay between a viral factor Nsp1 and the host translation factor EIF1A, selectively regulating CoV2 gRNA translation initiation in favor of coronavirus replication. This interdependency therefore presents a potential new target for antiviral therapies.

## Acknowledgments

This work was supported by NIH grants AI171421, AI137471, AI169460 to R.An. and NIH grants AI127447 and AI127447 to JF. We thank Angela Detweiler, Sheryl Paul, Honey Mekonen and Norma Neff of the Chan Zuckerberg Biohub San Francisco for their help with sequencing the ribo-seq libraries.

## Author contributions

R.Av., R.An. and J.F. conceived and coordinated the study. R.Av. designed and performed experiments, analyzed data and performed statistical analyses. P.L. and Y.X. performed all BSL3 experiments with live SARS-CoV-2. M.T. performed cloning. L.Z., P.L.M., and J.E. performed LC-MS/MS sample prep and data acquisition. R.Av. and R.An. wrote the manuscript.

## Data availability

Sequencing data were deposited in SRA database under BioProject number PRJNA932822. The mass spectrometry proteomics data have been deposited to the ProteomeXchange Consortium via the PRIDE partner repository with the dataset identifier PXD039981.

## PRIDE reviewer account details

eztwNtVI

## Figures

**Extended Data Figure 1.**
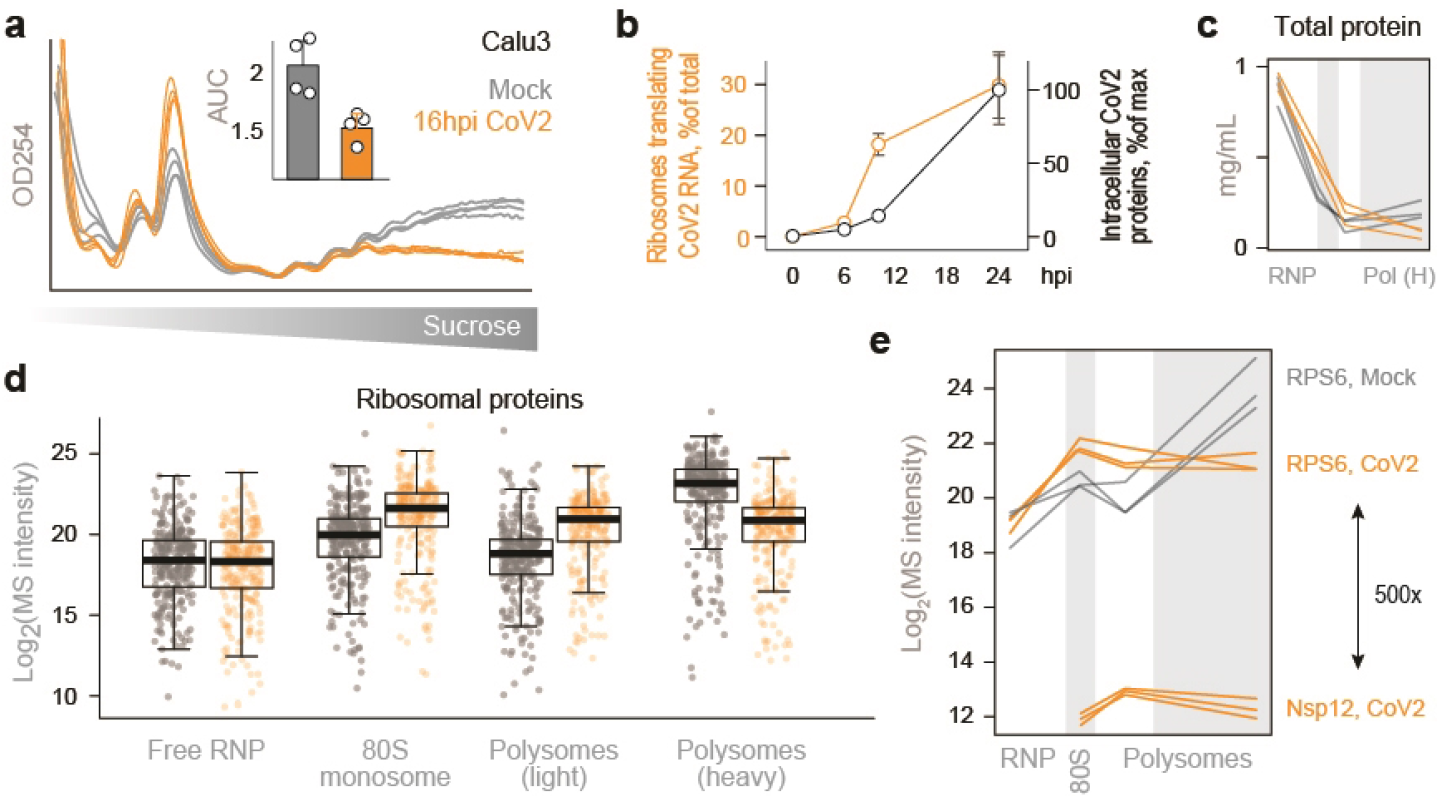
Subcellular proteomics of CoV2 infected cells. (**a**) Calu3 cells were infected with SARS-CoV-2 USA/WA1/2020 at MOI=5, lysed, fixed with formaldehyde and fractionated on 10-50% sucrose gradients with continuous monitoring of rRNA absorbance. Each line reflects a single replicate, and bar graphs show the ratio of polysomes to sub-polysomes, calculated as the area under the curve (AUC) of relevant fractions. Shown are means±SD of 4 independent replicates. (**b**) Timecourse of CoV2 RNA translation and intracellular viral protein accumulation, from^11^. Shown are means±SD of 3 independent replicates. (**c**) Total protein extracted from pooled fractions of infected and uninfected cells, quantified by bicinchoninic acid (BCA) assays. Each line reflects a single replicate. (**d**) Boxplots of all cytosolic ribosomal proteins quantified by MS in each pooled fraction from all 3 replicates. (**e**) Line plots of RPS6 and CoV2 Nsp12 (RdRp) quantified by MS in each pooled fraction. Each line represents a single replicate.

**Extended Data Figure 2.**
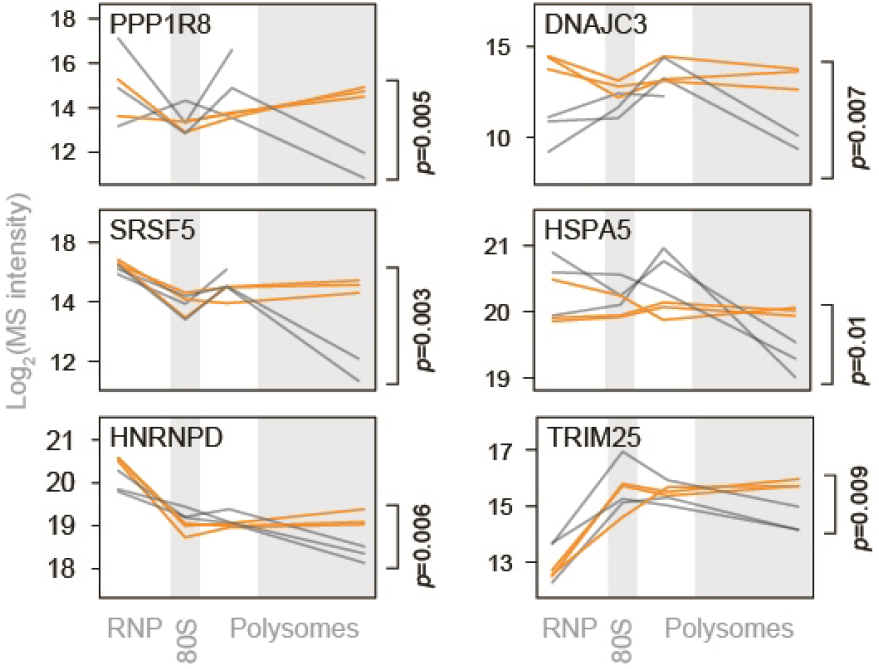
Proteostasis factors recruited to heavy polysome fractions upon CoV2 infection. Line plots of individual ribosomal proteins quantified by MS in each pooled fraction. Each line represents a single replicate. P, two-tailed Student’s t-test p-value of differences in indicated protein abundance in heavy polysome fractions.

**Extended Data Figure 3.**
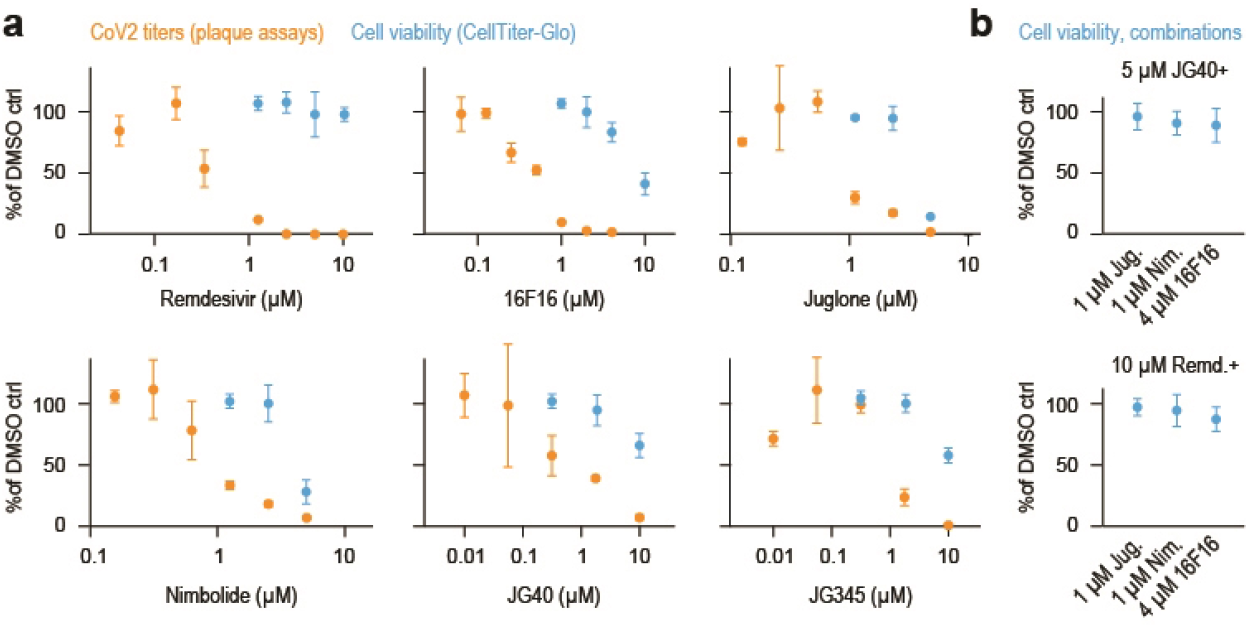
Toxicity and antiviral effects of proteostasis modulators. (**a-b**) Vero cells were infected with CoV2 at MOI=0.5. Single drugs (a) or drug combinations (b) were added at the start of infection, and titers were determined by plaque assays at 16 hours post-infection. Toxicity was determined using CellTiter-Glo at 24h of drug treatment, in the absence of CoV2 infection. Shown are means±SD of 3 independent replicates, normalized to DMSO controls.

**Extended Data Figure 4.**
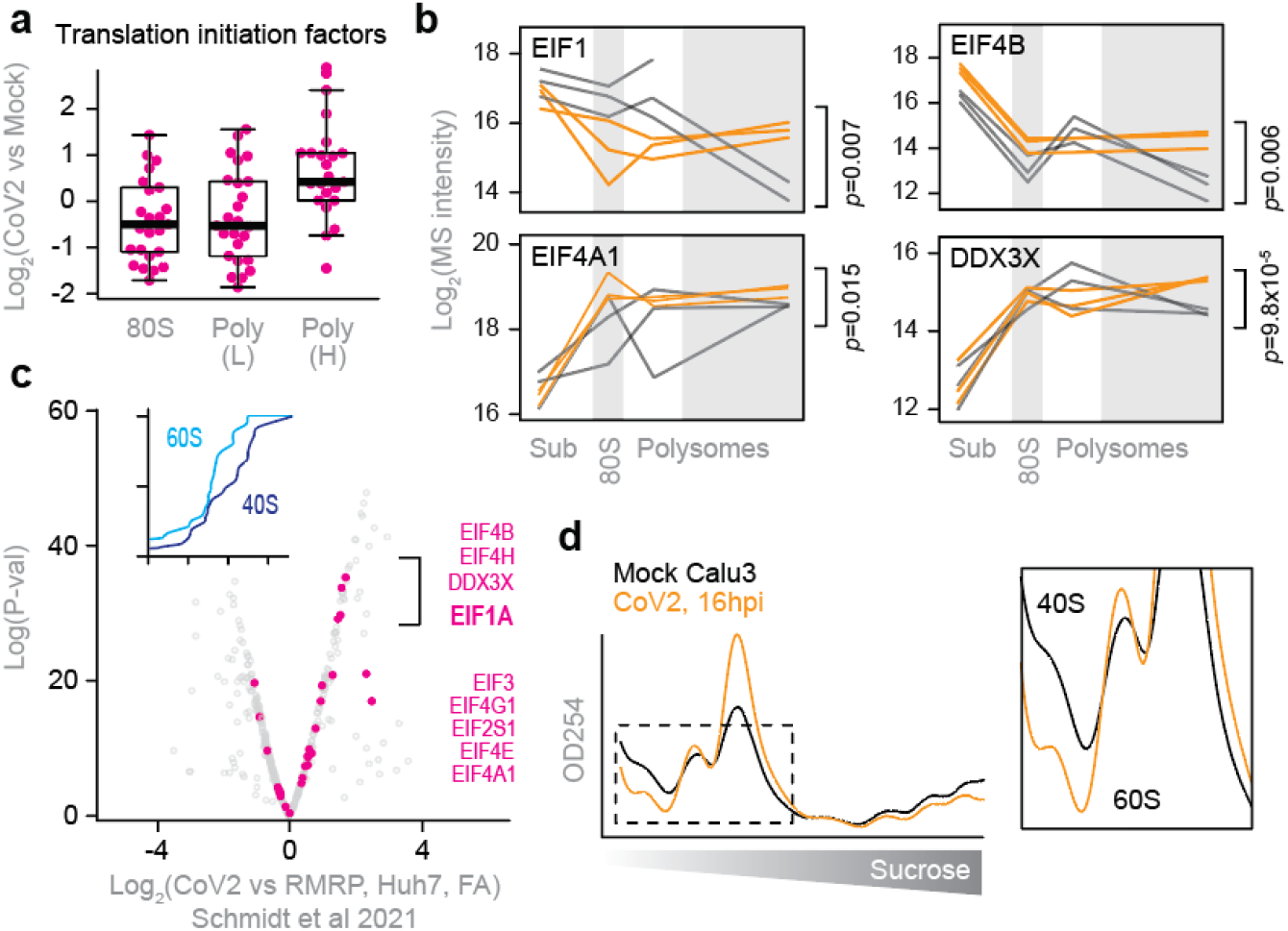
Inefficient translation initiation on CoV2 gRNA. (**a**) Change in abundance of individual translation initiation factors upon CoV2 infection, in each fraction. (**b**) Line plots of individual translation factors quantified by MS in each fraction. Each line represents a single replicate. P, two-tailed Student’s t-test p-value. (**c**) CoV2 RNA interactome is enriched for translation initiation factors during infection. Shown are pairwise comparisons of individual host protein abundance, quantified by MS, that specifically interact with either genomic or subgenomic CoV2 RNA. Inset, cumulative distribution plots of 40S and 60S ribosomal protein interaction with CoV2 RNA. (**d**) rRNA absorbance profiles showing lower abundance of free 40S subunits during CoV2 infection. Sum of four replicates.

**Extended Data Figure 5.**
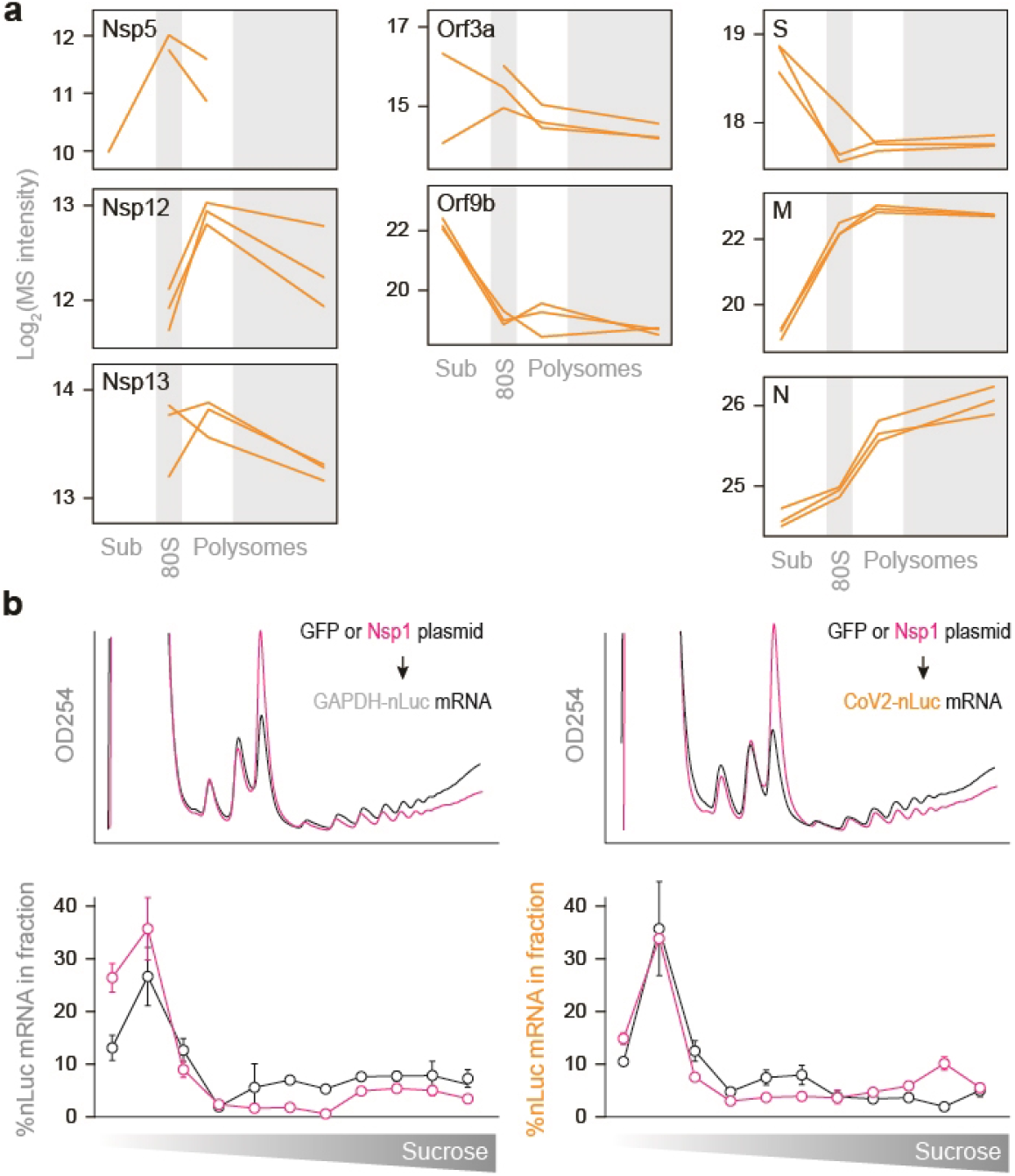
Sucrose sedimentation patterns for individual CoV2 proteins. (**a**) Line plots of individual viral proteins quantified by MS in each fraction. Each line represents a single replicate. (**b**) Related to Fig. 5d: Cells transfected with either GFP or Nsp1 followed by GAPDH-nLuc or CoV2-nLuc mRNA were lysed and fractionated on 10-50% sucrose gradients with continuous monitoring of rRNA. The content of nLuc mRNA in each fraction was determined by qPCR. Line plots show the proportion of nLuc mRNA in each fraction of a single gradient. Shown are means±SD of 2 independent replicates.

## Methods

### Cell cultures and viral infection

The African green monkey kidney Vero E6 (ATCC, no. CRL-1586) and human Calu3 cells (ATCC, no. HTB-55) were grown in DMEM medium (Thermo Fisher) supplemented with 10% fetal bovine serum, 100 units/ml penicillin and 100 mg/ml streptomycin, at 37°C in a 5% CO2 incubator. A clinical isolate of SARS-CoV-2 (USA-WA1/2020, BEI Cat No: NR-52281) was propagated in Vero E6 cells and was used for Vero E6 experiments. Viral titers were quantified with a plaque assay. All the infections were performed at biosafety level-3 (BSL-3). To assess the antiviral activity, ∼70% confluent monolayers of Vero E6 cells (3×105 cells/well in 24-well plates) were pretreated with drugs or drug combinations at indicated concentrations for 3 hours (pretreatment) and then infected with SARS-CoV-2 (MOI = 0.5) at 37°C for 1 hour. The virus solution was removed, cells were further cultured with fresh medium containing drugs at the same concentrations. At 16 hours post-infection, supernatants were collected, and viral titers were measured with a plaque assay.

### Plaque assay

Confluent monolayers of Vero E6 cells grown in six-well plates were incubated with the serial dilutions of virus samples (250 μl/well) at 37°C for 1 hour. Next, the cells were overlayed with 1% agarose (Invitrogen) prepared with MEM supplemented with 2% FBS and antibiotics. Three days later, cells were fixed with 4% formaldehyde for 2 hours, the overlay was discarded, and samples were stained with crystal violet dye, and the number of plaques was calculated.

### Polysome profiles

A total of 1-5×10^7^ Vero E6 or Calu3 cells in T175 flasks were washed twice with and scraped into ice-cold PBS with calcium and magnesium. Cells were pelleted at 1,200 RCF for 5 min at 4°C, and supernatants were removed. Pellets were resuspended in 220 µl polysome buffer (25 mM HEPES pH=7.5, 100 mM NaCl, 10 mM MgCl_2_, 2 mM dithiothreitol (DTT) and Complete EDTA-free protease inhibitor cocktail (Millipore Sigma). Triton X-100 and sodium deoxycholate were added to a final concentration of 1% each and the samples were incubated on ice for 20 min and centrifuged at 20,000 RCF for 10 min at 4°C to remove cell debris. 250 µl lysates were transferred to fresh tubes, combined with 125 µl 12% formaldehyde, and incubated on ice for 30 min. To quench the crosslinking reaction, samples were incubated with 125 µl 4M Tris pH=8.0 on ice for 20 min, followed by flash freezing. Frozen lysates were thawed on ice and loaded on 10-50% sucrose gradients in Polysome buffer and subjected to ultracentrifugation at 36,000 rpm in an SW41.Ti swinging bucket rotor (Beckman Coulter) for 150 min at 4°C. 15 Equal volume fractions were collected using Gradient Station (BioComp) with continuous monitoring of rRNA at UV254. Fractions were pooled as follows: 1-4 for free RNP; 5 for 80S monosomes; 6-8 for light polysomes, and 9-15 for heavy polysomes. For polysome profiles of cells transfected with nanoluciferase reporters, the above protocol was followed without formaldehyde fixation or pooling of gradient fractions.

### Sample preparation for proteomic analysis

Pooled gradient fractions were diluted 1:2 with ice-cold PBS and tumbled overnight at 4°C with 50 µl Strataclean resin (Agilent). Beads were pelleted at 600 RCF for 5 min at 4°C, and supernatant was removed. Beads were resuspended in 100 µl PBS supplemented with 2% SDS and incubated at 95°C for 15 min to elute proteins from beads and reverse formaldehyde crosslinks. Beads were pelleted at 600 RCF for 5 min at room temperature, and supernatants were transferred to fresh tubes. Protein was extracted using methanol-chloroform precipitation: 400 µl methanol, 100 µl chloroform and 350 µl water were added sequentially to each 100 µl sample, followed by centrifugation at 14,000 RCF for 5 min at room temperature. The top phase was removed, and the protein interphase was precipitated by addition of 400 µl methanol, followed by centrifugation at 14,000 g for 5 min at room temperature. Pellets were air dried and resuspended in 8M urea, 25 mM ammonium bicarbonate (pH 7.5). Protein concentration was determined by BCA (Thermo Fisher) and 1-2 µg total protein were subjected to reduction and alkylation by incubation with 10 mM DTT for 1 h at room temperature followed by 5 mM iodoacetamide for 45 min at room temperature, in the dark. The samples were then incubated with 1:50 enzyme to protein ratio of sequencing-grade trypsin (Promega) overnight at 37 °C. Peptides were desalted with μC18 Ziptips (Millipore Sigma), dried and resuspended in 10 μL 0.1% formic acid in water.

### LC-MS/MS acquisition

Digested peptides were resuspended in 0.1% formic acid and 40% of each specimen was analyzed on an LTQ Orbitrap Fusion Lumos Tribrid Mass Spectrometer (Thermo Fisher Scientific) coupled with a Dionex Ultimate 3000 liquid chromatography system (Thermo Fisher Scientific). Peptides were separated by capillary reverse-phase chromatography for 120 min on a 24-cm reversed-phase column (inner diameter of 100 μm, packed in-house with ReproSil-Pur C18-AQ 3.0 m resin (Dr. Maisch)). A multi-step linear gradient was applied as follows: 96% A + 4% B to 75% A + 25% B over 70 min; 75% A + 25% B to 60% A + 40% B over 20 min; 60% A + 40% B to 2% A + 98% B over 2 min and maintain this proportion for 2 min before returning to 98% A + 2% B and holding this proportion for 24 min. Buffer A is 0.1% formic acid in water and buffer B is 0.1% formic acid in acetonitrile; flow rates were maintained at 300 nl/min throughout the gradient. Full MS scans of intact peptide precursor ions were acquired in the Orbitrap mass analyser at a resolution of 120,000 (FWHM) and m/z scan ranges of 400–1,500 in a data-dependent mode. The automatic gain control (AGC) target was 4 × 10^5, the maximum injection time was 50 ms, and the isolation width was 1.6 Da. The most intense ions recorded in full MS scans were then selected for MS2 fragmentation in the Orbitrap mass analyser using higher-energy collisional dissociation (HCD) with a normalized collision energy of 30% and resolution of 15,000 (FWHM). Monoisotopic precursor selection was enabled, and singly charged ion species and ions with no unassigned charge states were excluded from MS2 analysis. Dynamic exclusion was enabled, preventing repeated MS2 acquisitions of precursor ions within a 10 ppm m/z window for 15 s. AGC targets were 5 × 10^4 and the maximum injection time was 100 ms.

### Mass spectrometry data processing

Raw MS data were processed using MaxQuant version 1.6.7.0 ^38^. MS/MS spectra searches were performed using the Andromeda search engine^39^ against the forward and reverse human and mouse Uniprot databases (downloaded August 28, 2017 and November 25, 2020, respectively). Cysteine carbamidomethylation was chosen as fixed modification and methionine oxidation and N-terminal acetylation as variable modifications. Parent peptides and fragment ions were searched with maximal mass deviation of 6 and 20 ppm, respectively. Mass recalibration was performed with a window of 20 ppm. Maximum allowed false discovery rate (FDR) was <0.01 at both the peptide and protein levels, based on a standard target-decoy database approach. The “calculate peak properties” and “match between runs” options were enabled. All statistical tests were performed with Perseus version 2.0.7.0 using either ProteinGroups or Peptides output tables from MaxQuant. Potential contaminants, proteins identified in the reverse dataset and proteins only identified by site were filtered out. For proteins not annotated in the Chlorocebus sabaeus proteome, we manually added the human ortholog gene symbol where similarity was >90%. Human ortholog Uniprot IDs were added based on gene symbol. Intensity-based absolute quantification (iBAQ) was used to estimate absolute protein abundance. Data was log2 transformed and median adjusted. Two-sided Student’s t-test with a permutation-based FDR of 0.01 and S0 of 0.1 with 250 randomizations was used to determine statistically significant differences between grouped replicates. Categorical annotation was based on Gene Ontology Biological Process (GOBP), Molecular Function (GOMF) and Cellular Component (GOCC), as well as protein complex assembly by CORUM.

### Drug treatments, toxicity and synergy

To determine the effects of drugs on host cells, 5×10^4^ Vero E6 cells were seeded per well in a black wall, clear bottom 96-well plate. Drugs were added 24 h after seeding. 24 h after adding drugs, cell viability was assayed using CellTiter-Glo (Promega), according to the manufacturer’s instructions, on a Tecan Ultra Evolution microplate reader. To determine the effects of drugs on CoV2 infection, virus adsorption was performed as described above, at MOI=0.5, and inoculum was replaced with fresh media containing drugs. Cell supernatants were collected at 16 hpi and subjected to plaque assays. SynergyFinder2^40^ was used to calculate Bliss scores.

### Nanoluciferase reporters, viral proteins, and RNA/DNA transfections

Plasmids encoding for nanoluciferase (nLuc) flanked by 5’ and 3’ untranslated regions of either CoV2 or GAPDH, and plasmids encoding for strep-tagged individual CoV2 proteins were kind gifts from Joseph (Jody) Puglisi^10^ and Nevan Krogan^4^. To generate translation-competent mRNA, nLuc plasmids were linearized using SpeI and subjected to in-vitro transcription using HiScribe T7 kit (New England Biolabs), according to the manufacturer’s instructions. In 20 μL total volume per reaction, 1 μg plasmid was combined with 2 μL buffer, 2 μL of each nucleotide, 1.6 μL cap, 1 μL SUPERase-in and 2 μL enzyme. Transcription was performed for 3 h at 37°C and terminated by precipitation. Each sample was combined with 1:1 v/v 5 M ammonium acetate, incubated on ice for 15 min, and centrifuged 20,000 RCF for 30 min at 4°C. Pellets were washed with 1 mL 75% ethanol, air dried and resuspended in 50 μL nuclease-free water. RNA integrity was confirmed using agarose gels stained with SYBR Safe, and concentration was determined by nanodrop.

For RNA transfections, Vero E6 cells were plated at 2×10^4^ cells/well in 96-well plates. The following day, transfection reactions were prepared as follows (per well). 0.15 μL Lipofectamine MessengerMAX (Thermo) was diluted in 5 μL OptiMEM (Thermo) and incubated for 10 min at RT. 20 ng in-vitro transcribed RNA was diluted in 5 μL OptiMEM and mixed with the MessengerMAX solution for 5 min. 10 μL transfection reactions were added per well, and cells were harvested at 4 h post-transfection. Luminescence measurements were acquired using Nano-Glo Luciferase Assay System (Promega), according to the manufacturer’s instructions, on a Tecan Ultra Evolution microplate reader.

For DNA/RNA transfections, Vero E6 cells were plated at 2×10^4^ cells/well in 96-well plates. The following day, transfection reactions were prepared as follows (per well). 0.1 μL Lipofectamine 2000 (Thermo) was diluted in 5 μL OptiMEM and 2 ng plasmid DNA was diluted in 5 μL OptiMEM. The two solutions were mixed, incubated at RT for 5 min, and added to cells. Media was replaced at 6 h post-transfection. RNA transfections were performed as above, 24 h after DNA transfections. For DNA/RNA transfections prior to sucrose gradient fraction, the above procedure was scaled up 500x, and cells were collected for polysome profiled analysis, as described above.

### Quantitative Real-Time PCR (qRT-PCR)

Total RNA was extracted using Trizol (Invitrogen) according to the manufacturer’s instructions. To extract RNA from sucrose gradient fractions, 1 µl pellet paint co-precipitant (Millipore Sigma) and 500 µl phenol:chloroform:isoamyl alcohol (25:24:1) were added to each 500 µl fraction and incubated for 5 min at room temperature. Phase separation was performed at 12,000 RCF for 15 min at 4°C, and the top phase was removed and subjected to another round of extraction as above. 400 µl of the top phase was combined with 600 µl isopropanol and RNA was pelleted at 12,000 RCF for 1 h at 4°C. Pellets were washed with 1 ml 75% ice-cold ethanol, air dried and resuspend in 20 µl RNase-free water. cDNA was synthesized using the High Capacity cDNA Reverse Transcription Kit (Thermo Fisher), according to the manufacturer’s instructions, using 5 µl of RNA from each gradient fraction. qRT-PCR analysis was performed using SensiFast SYBR (BioLine) and gene-specific primers (nLuc FWD 5’ CAGCCGGCTACAACCTGGAC 3’, REV 5’ AGCCCATTTTCACCGCTCAG 3’; GAPDH FWD 5’ AGGTCGGAGTCAACGGAT 3’, REV 5’ TCCTGGAAGATGGTGATG 3’), according to the manufacturer’s instructions. To estimate relative abundance of specific mRNAs in each gradient fraction, each Ct value was divided by the sum of Ct values across all gradient fractions.

### siRNA transfections

siRNAs against EIF1A (Dharmacon Smartpool M-011262-02-0005 and M-011908-00-0005, targeting both EIF1AX and EIF1AY), as well as nontargeting siRNAs (Smartpool D-001206-13-05), were from Dharmacon. siRNAs were reconstituted in nuclease-free water to a concentration of 10 µM. Vero E6 cells were plated at 1×10^5^ cells/well of 12-well plates. The following day, 3 µl RNAiMax reagent (Thermo) was diluted in 50 µl OptiMEM and combined with 2 µl siRNA in 50 µl OptiMEM. For EIF1A, 1 µl of each siRNA pool was used. After 5 min at RT, the reaction mix was added to cells, and media was replaced after 6 h. At 48 h post-transfection, cells were passaged and transfected again, as above. Additional DNA/RNA transfections were launched 48 h after the second siRNA transfection.

### SDS-PAGE and immunoblotting

For immunoblotting, cells were washed twice with ice-cold PBS and lysed on plate with RIPA buffer (25 mM Tris-HCl pH=7.5, 150 mM NaCl, 1% NP-40, 0.5% Sodium deoxycholate) supplemented with 2 mM DTT, Complete EDTA-free protease inhibitor cocktail, and 50 units/mL benzonase (Millipore Sigma) to remove DNA. Lysis was performed on ice for 20 min and lysates were clarified by centrifugation for 10 min at 12,000 RCF at 4°C. Protein concentration was determined by BCA assay (Thermo Fisher) and 4x Laemmli sample buffer (Bio-Rad) supplemented with fresh 10% 2-mercaptoethanol was added to a final concentration of x1. 15 µg of each sample was resolved on 4-20% SDS-PAGE (Bio-Rad), transferred to 0.2 µm PVDF membranes presoaked in methanol for 30 sec. Membranes were blocked with 4% molecular biology grade BSA (Millipore Sigma) in tris-buffered saline supplemented with 0.1% Tween-20 (Millipore Sigma) (TBST) for 1 h at room temperature then probed with specific primary antibodies for 2 h at room temperature. Primary antibodies were diluted in 4% BSA/TBST supplemented with 0.02% sodium azide, as follows: rabbit anti-EIF1A (1:1000, GeneTex GTX118810), rabbit anti-EIF1 (1:1000, ProteinTech 15276-1-AP), mouse anti-beta tubulin (1:10,000, EMD 05-661). Secondary antibodies were diluted 1:10,000 in TBST. Western blot detection was done using ECL Plus Western Blotting Substrate (Thermo Fisher) and images were taken either by film radiography.

### Ribosome profiling analysis

Ribosome footprints were prepared essentially as described^41^. EIF1A and nontargeting siRNA were transfected into Vero E6 cells as above, and 24 h after the second transfection, cells were replated at 2×10^6^ per T25 flask. CoV2 infection was launched the following day at MOI=5. At 16 hpi, cells were washed with 5 mL ice-cold PBS with calcium and magnesium. PBS was fully aspirated and replaced with 350 µl polysome buffer (25 mM HEPES pH=7.5, 100 mM NaCl, 10 mM MgCl_2_, 2 mM DTT and Complete EDTA-free protease inhibitor cocktail) supplemented with 1% Triton X-100 and 1% sodium deoxycholate. On-plate lysis was allowed to proceed on ice for 10 min with occasional shaking. Lysates were transferred to fresh tubes and clarified at 12,000 RCF for 5 min at 4°C. 200 µl clarified lysate were combined with 5 µl RNase I (10 U/µl, Invitrogen). Reactions were incubated at RT for 45 min with shaking, and terminated by addition of 200 U Superase-In (Thermo). Spin columns were used to separate monosome, as follows. S-400 HR MicroSpin Columns (Amersham) were drained at 600 RCF for 4 min at 4°C and the flowthrough was discarded. Resin was resuspended in 200 µl polysome buffer and centrifuged again. Each 200 µl RNase-digested sample was loaded on 2 separate columns (100 µl each), centrifuged at 600 RCF for 2 min at 4°C. The flowthroughs were combined and 800 µl Trizol (Invitrogen) was added. RNA was extracted according to the manufacturer’s instructions. Footprints were resolved on 8M urea 15% PAGE, according to the protocol^41^. 15-35 nt fragments were extracted, dephosphorylated, linker ligated according to the protocol. rRNA depletion was performed using Ribo-Zero rRNA Removal Kit (Illumina), according to the manufacturer’s instructions. All downstream steps were performed as described^41^. Libraries from two independent repeats were sequenced on a NextSeq 2000 P3 (Illumina). After demultiplexing, sequencing reads were trimmed of adaptor sequences and quality filtered using cutadapt (-a CTGTAGGCACCATCAAT -m1 -q20). Ribosomal RNA was removed using Bowtie2 (--un). Remaining reads were aligned to SARS-Cov-2 genomes [Genebank MN985325] using Hisat2 (--trim5 1). Per-nucleotide ribosome occupancy tables were generated using bamCoverage and visualized on Integrated Genome Viewer.

